# Monosynaptic tracing success depends critically on helper virus concentrations

**DOI:** 10.1101/736017

**Authors:** Thomas K. Lavin, Lei Jin, Nicholas E. Lea, Ian R. Wickersham

## Abstract

Monosynaptically-restricted transsynaptic tracing using deletion-mutant rabies virus has become a widely used technique in neuroscience, allowing identification, imaging, and manipulation of neurons directly presynaptic to a starting neuronal population. Its most common implementation is to use Cre mouse lines in combination with Cre-dependent “helper” adeno-associated viral vectors (AAVs) to supply the required genes to the targeted population before subsequent injection of a first-generation (ΔG) rabies viral vector. Here we show that the efficiency of transsynaptic spread and the degree of nonspecific labeling in wild-type control animals depend strongly on the concentrations of these helper AAVs. Our results suggest practical guidelines for achieving good results.

## INTRODUCTION

Rabies virus has proven useful for neuroscience, because of its natural behavior of spreading between synaptically-connected neurons (although both the mechanism of transsynaptic spread and the true degree of its synaptic specificity remain unclear[1; 2]) in an apparently exclusively retrograde direction (in the central nervous system, whereas in primary sensory neurons it appears to be bidirectional[3; 4]). This has allowed it to serve as a useful tool for mapping synaptic connections, usually in the context of “monosynaptic tracing”, which refers to the use of a modified rabies virus to label neurons that are, putatively, directly presynaptic to a targeted population of neurons[5]. It relies on the use of a rabies virus to which two modifications have been made. First, in order to render it incapable of spread between neurons without assistance, one (or more) of its genes has been deleted; in first-generation vectors, this is the glycoprotein gene, denoted “G”[6]. Second, in order to allow selective targeting of the initial infection to the group of neurons of interest, the viral particles are coated with the envelope protein of a different virus (the avian-endemic retrovirus “ASLV-Awhich refers to the use of a modified rab; the envelope protein of which is called “EnvA”), rendering the virus incapable of infecting mammalian neurons without assistance[5]. In the targeted neuronal population, two exogenous proteins must be expressed before injection of the rabies virus: the receptor for EnvA (a quail cell surface protein called “TVA”[7]), to allow the modified rabies virus to infect the starting cells, and the deleted viral gene(s) (G, in the case of first-generation (“ΔG”) vectors). While this can be achieved by single-cell transfection techniques[8; 9; 10; 11], the much more accessible and widely used implementation is to use Cre[12]-dependent adeno-associated viral vectors (“AAVs”)[13; 14; 15; 16; 17; 18; 19; 20; 21] in combination with a Cre mouse line, in order to map inputs to a Cre-expressing group of neurons (see Figure 1). This approach has been used in a large number of studies and contributed considerably to our understanding of the organization of many circuits within the mammalian nervous system.

**Figure 1:**
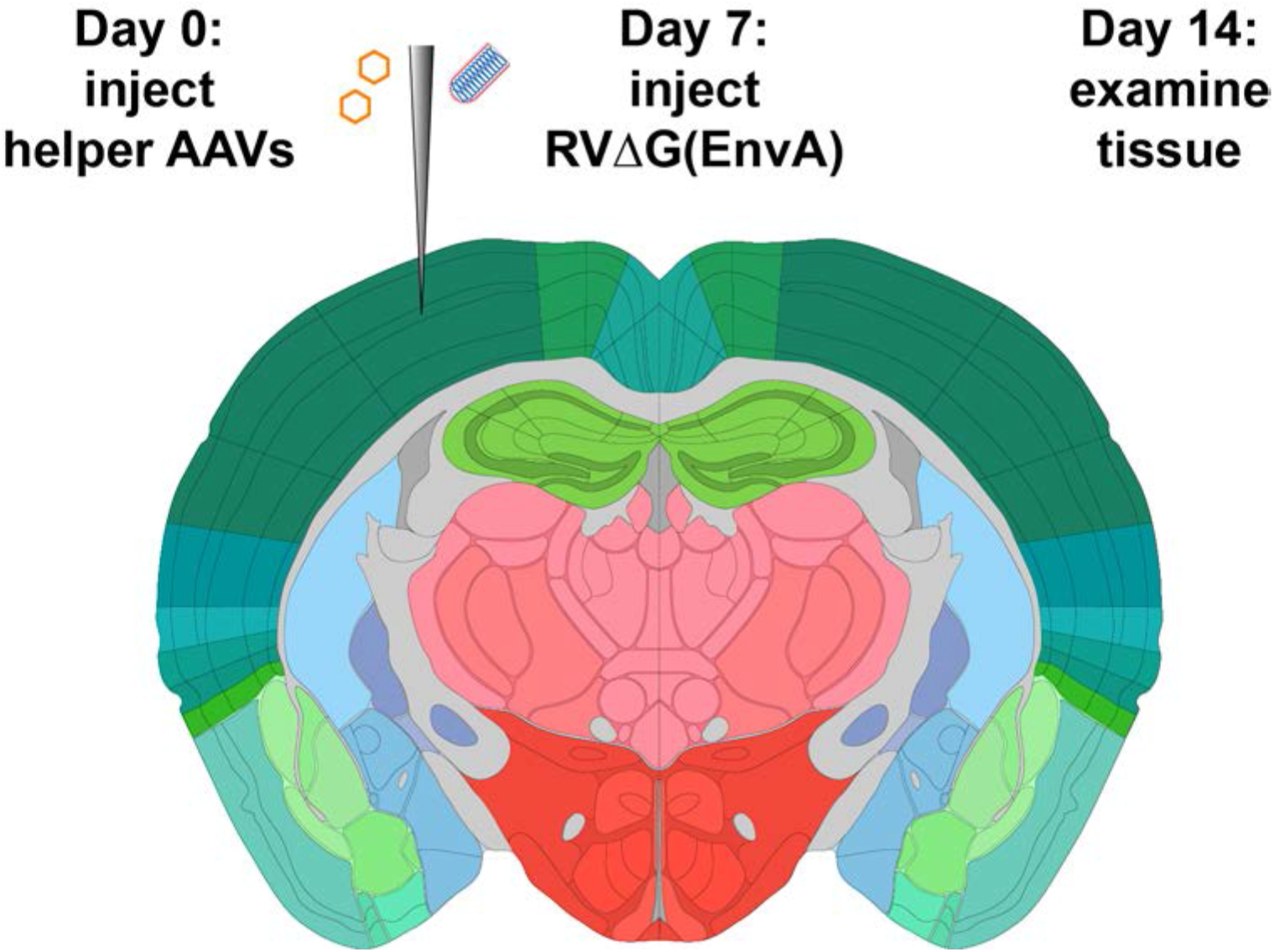
Strategy for monosynaptic tracing with helper AAVs. Helper viruses are injected in a Cre mouse, or a Cre-negative mouse in the case of control experiments, then rabies virus (RV) (ΔG, EnvA-coated, expressing mCherry) is injected in the same location subsequently. While different labs have used various intervals and survival times, we used a 7-day interval between AAV and RV injections, and another 7-day interval between RV injection and perfusion, in all experiments for this paper.

Within this basic paradigm of using Cre-dependent “helper” AAVs to provide the genes required for monosynaptic tracing, many variations are possible. A major consideration in design of such experiments is the mismatch between the miniscule amount of TVA required for successful initial infection of the starting cells, because of the high sensitivity of the EnvA-TVA interaction[22; 23], and the high levels of G that appear to be required for efficient spread of the virus from the starting cells to the putatively presynaptic cells. This causes a problem: to achieve acceptable levels of transsynaptic spread, a high concentration of AAV may be found to be necessary; however, because it appears that all putatively Cre-dependent AAVs “leak” because of spontaneous recombination of some fraction of virions’ genomes, even at the plasmid DNA stage (unpublished results and Kimberly Ritola, personal communication), a high concentration of an AAV expressing TVA can result in an unacceptably high level of “background” labeling of cells by the rabies virus in wild-type mice. Different groups have taken different approaches to dealing with this, including using two separate Cre-dependent AAVs so that G is expressed at a presumably higher level than TVA[15] and/or using a low-affinity mutant of TVA[24; 25].

Our current approach is to use a mixture of two AAVs, described first in Liu et al.[18], which are coupled by the tetracycline transactivator system[26]. The first AAV is Cre-dependent and expresses, in Cre-expressing cells, TVA (transmembrane isoform[7; 27; 28]), EGFP[29], and the tetracycline transactivator (“tTA”); the second is not directly Cre-dependent but simply expresses both G (SAD B19 strain[30]) and the blue fluorophore mTagBFP2[31] under the control of the tetracycline response element. Expression of tTA from the first AAV drives expression of G from the second AAV in the same cells. Note that this uses the “TET-OFF” system, so that no tetracycline (or doxycycline, etc.) needs to be added in order to make the system work.

The use of this combination has several intended advantages. First, the use of a two-AAV combination allows the concentrations of the vectors encoding TVA and G to be independently titrated. Second, the tTA-TRE system should provide amplification of the level of G expression with respect to the level of expression of the genes in the first AAV, so that the TVA/EGFP/tTA virus can be titrated down to very low concentration to result in low background labeling in wild-type mice but with the G expression level still high enough to result in plentiful transsynaptic spread of the rabies virus. Third, the use of the tet transactivator system can also allow the expression of G (and mTagBFP2) to be turned off (or potentially titrated) by administration of doxycycline after transsynaptic spread has taken place, in order to mitigate toxicity to the starting cells, although we have not done this in any of the work presented in this paper.

We have recently published a detailed step-by-step protocol for monosynaptic tracing using these viruses for monosynaptic tracing with Cre mice[32]. Here we present results of our titration experiments to test the effects of using different dilutions of the helper viruses, to show the reasons for the specific concentrations that we recommend. We found that the two-helper combination described above and in Liu et al.[18] gives much better results than the simpler single AAV which we described earlier[16], which did not allow independent optimization of transsynaptic tracing efficiency and minimization of background labeling in Cre-negative mice. We also found that *excessively high* titers of the helper viruses gave very poor results, suggesting that preventing toxicity due to overly high expression of the helper virus genes is as important as ensuring sufficient expression of them. Finally, and most practically, we suggest specific concentrations of the helper viruses that gave best results in the Cre line in which we performed the titration and that should serve either as a likely choice of parameters for end-users or as a promising starting point for much more limited titration series to be done as pilot experiments when targeting other cell types in other Cre lines.

## RESULTS

Having previously found Addgene’s Viral Service (www.addgene.org/viral-service/aav-prep) to be an excellent source of high-quality AAVs, we authorized them to package and distribute three of our published Cre-dependent helper AAVs: the standalone helper virus AAV-syn-FLEX-splitTVA-EGFP-B19G from Kohara et al.[16] (referred to below as the “tricistronic” helper virus) and the two viruses to be used in combination as described in Liu et al.[18]: AAV-syn-FLEX-splitTVA-EGFP-tTA and AAV-TREtight-mTagBFP2-B19G. Although the resulting preparations, all with serotype 1 capsids, had much higher titers than earlier batches that we had previously used successfully for similar experiments, we nonetheless first tried using them “straight”: undiluted except insofar as, for the two-helper-virus combination, the two viruses were combined in a 50/50 mixture by volume (see Methods). We injected either AAV1-syn-FLEX-splitTVA-EGFP-B19G or the two-virus mixture into the primary somatosensory cortex of either the PV-Cre (expressing Cre from the parvalbumin locus)[33] mouse line crossed with the Ai65F reporter line (FLP-dependent tdTomato[34])[35] or, for Cre-negative controls, the Ai65F reporter line without a Cre allele. 7 days after AAV injection, we injected RVΔG-4FLPo(EnvA)[32] (expressing FLPo recombinase[36]) and perfused the mice 7 days after that.

The results were surprisingly bad. Figure 2, panels a-h show example results using the two-helper combination without dilution. While many cells were well-labeled with EGFP, blue fluorescence was barely visible, and there were few tdTomato-labeled cells at the injection site and almost none elsewhere (see Figure 4 for quantification). Furthermore, matched control injections of the same viruses in Cre-negative mice resulted in undesirably large numbers of RV-labeled neurons at the injection site (Figure 2, panels i-l). This was evidently not the fault of the rabies virus preparation: in control animals injected only with rabies virus, with no helper viruses, we found very few RV-labeled cells at the injection site or otherwise (Supplementary Figure S1; cell counts are given in Supplementary Table S1). While results using the undiluted single tricistronic helper virus looked better in PV-Cre mice, they were still unimpressive, and the problem of label in Cre-negative mice was far worse (see Figure 4 for quantification; example images not shown in these cases).

**Figure 2:**
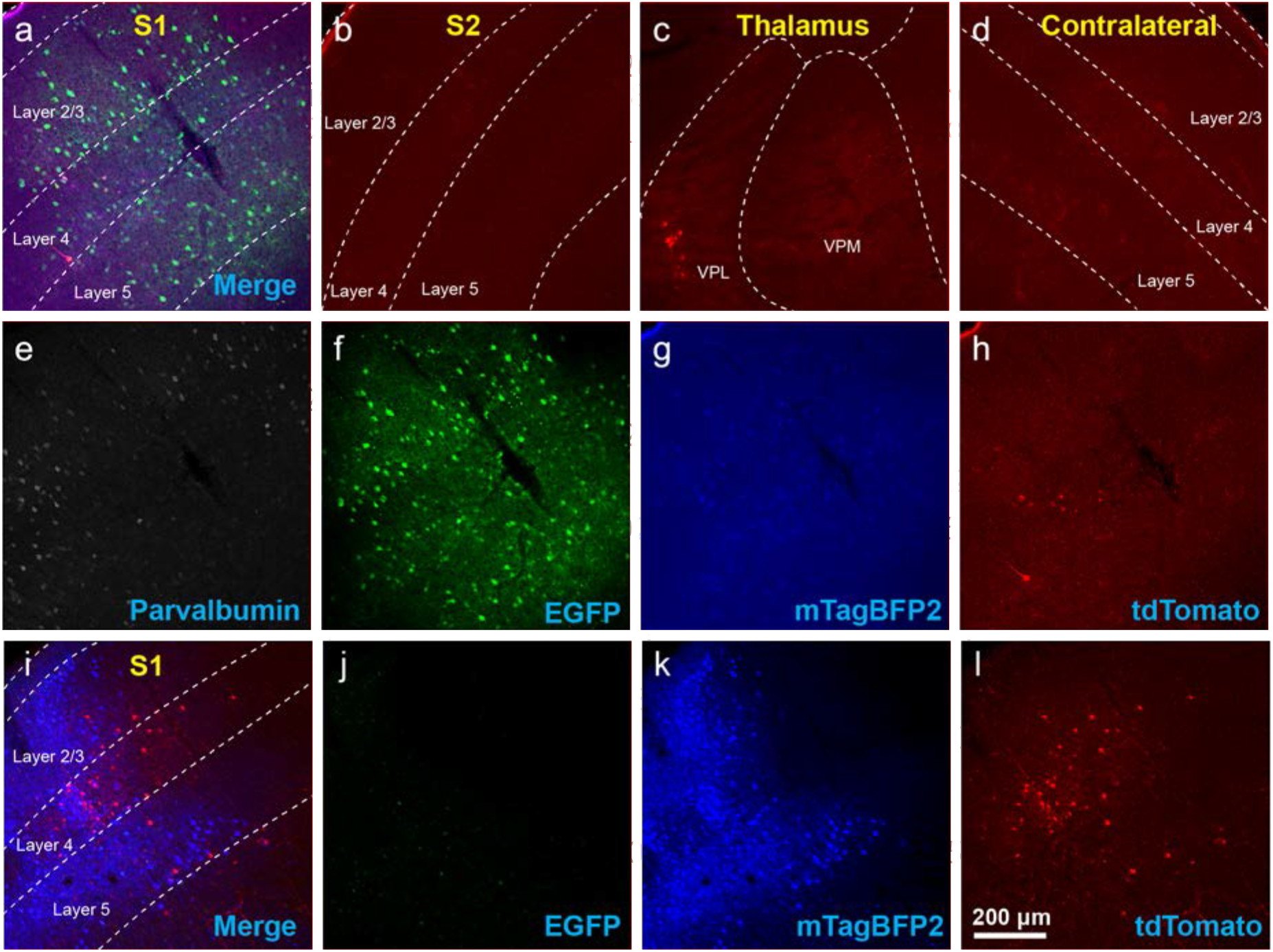
Use of helper viruses at excessive concentrations can result in near-complete failure of monosynaptic tracing. The two-AAV combination described in Liu et al.[18] (AAV1-synP-FLEX-splitTVA-EGFP-B19G mixed with AAV1-TREtight-mTagBFP2-B19G) was injected in somatosensory cortex of PV-Cre x Ai65F (FLPo-dependent tdTomato reporter) mice, followed by RVΔG-4FLPo(EnvA) 7 days later. a-h) Very poor results were obtained when using new preparations of these AAVs undiluted. a) Injection site in S1. Green = anti-EGFP staining, blue = mTagBFP2, red = tdTomato. Individual channels from this field are shown in panels e-h. b) No labeled neurons were found in ipsilateral secondary somatosensory cortex (S2). c) Very few labeled neurons were found in ipsilateral thalamus (VPL and VPM). d) No labeled neurons were found in contralateral S1. e-h) Individual channels from the field shown in panel a. e) anti-parvalbumin staining (not shown in panel a). f) anti-EGFP staining, indicating expression from the first, Cre-dependent AAV. g) mTagBFP2, indicating expression from the second, tTA-dependent AAV. h) tdTomato, reporting activity of the FLPo-encoding rabies virus. i-l) Injection site after using undiluted viruses in two-helper combination in Cre-negative animal: many labeled cells are seen. i) Overlay of j-l. j) anti-EGFP staining, k) mTagBFP2 signal, l) tdTomato marking RV labeling. Scale bar in a: 200 μm, applies to all panels.

On the assumption that the poor results in Cre mice were due to toxicity resulting from excessive concentration, we diluted the new preparations to the same titers as those of the batches that we had been using previously: namely, we diluted AAV1-syn-FLEX-splitTVA-EGFP-tTA by a factor of 17.96 (to 9.47 × 10^11 gc/ml, based on the titer reported by Addgene) and AAV1-TREtight-mTagBFP2-B19G by a factor of 19.98 (to 1.60 × 10^12 gc/ml, based on the titer reported by Addgene).

Using these diluted helper viruses gave us much better results, with large numbers of RV-labeled neurons found in upstream regions including secondary somatosensory cortex, thalamus, and contralateral cortex (Figure 3, panels a-h shows example results; see Figure 4 for quantification). However, in mice not expressing Cre, even using the diluted AAVs resulted in unreasonably-high numbers of RV-labeled neurons (red) (Figure 3, panels i-l, with quantification in Figure 4), as well as bright blue (but no visible green, even with immunostaining) labeling indicative of leaky expression from the helper viruses in the absence of recombinase.

**Figure 3:**
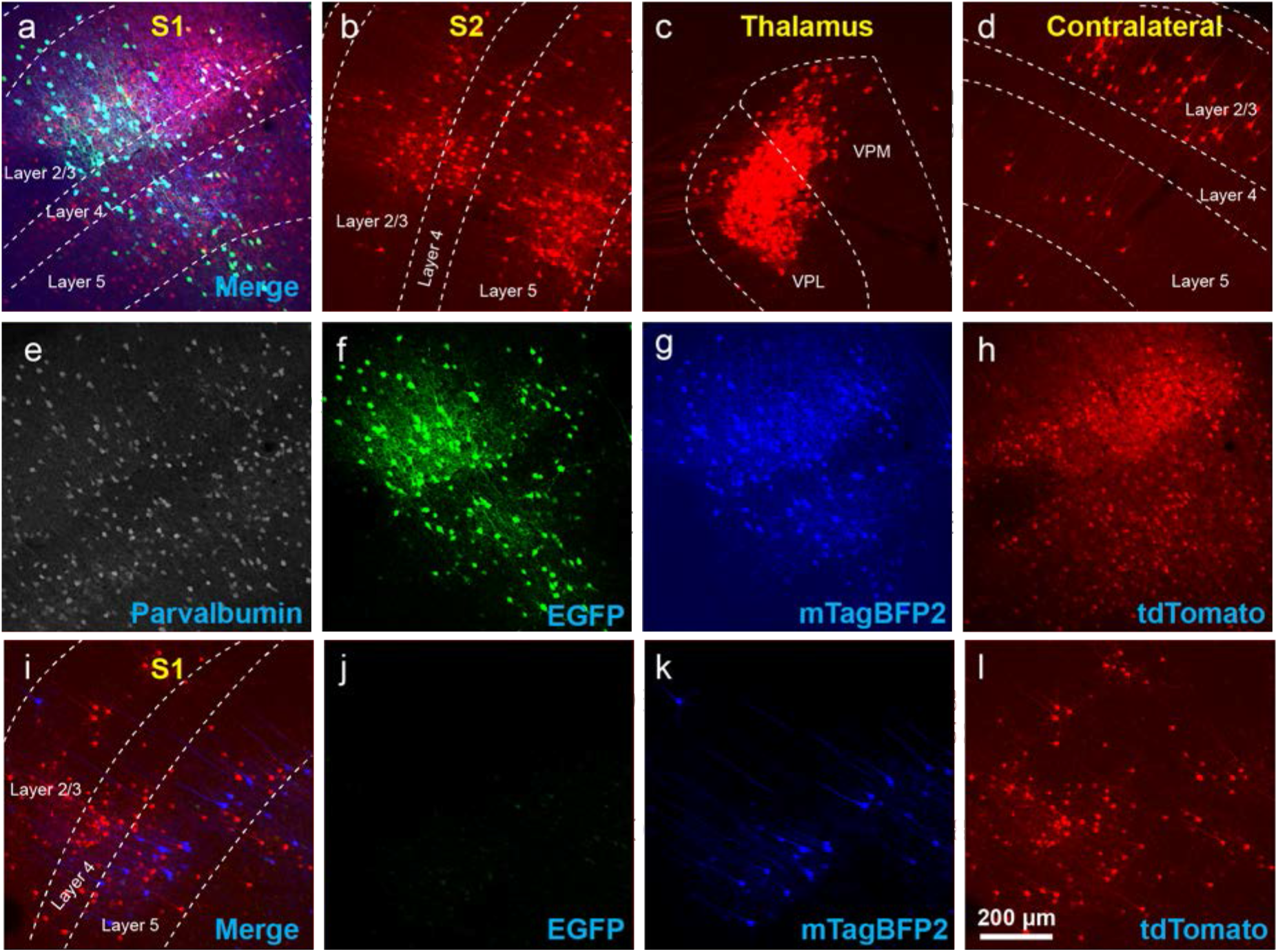
Use of insufficiently-diluted helper viruses results in excessive background labeling in Cre-negative animals. a-h) Diluting the helper viruses to concentrations matching previously used preparations gave much better results. a) Injection site in S1; individual channels from this field are shown in panels e-h. b-d) Many labeled presynaptic neurons were found in ipsilateral secondary somatosensory cortex (b), ipsilateral thalamus (VPL and VPM) (c), and contralateral S1 (d). e) anti-parvalbumin staining (not shown in panel a). f) anti-EGFP staining, indicating expression from the first, Cre-dependent AAV. g) mTagBFP2, indicating expression from the second, G-encoding AAV. h) tdTomato, reporting activity of the FLPo-encoding rabies virus. i-l) Even with the AAVs diluted to match the titers of previous batches, excessive background labeling is seen at the injection site. i) Overlay of j-l. j) anti-EGFP staining, k) mTagBFP2 signal, l) tdTomato marking RV labeling. Scale bar in a: 200 μm, applies to all panels.

**Figure 4:**
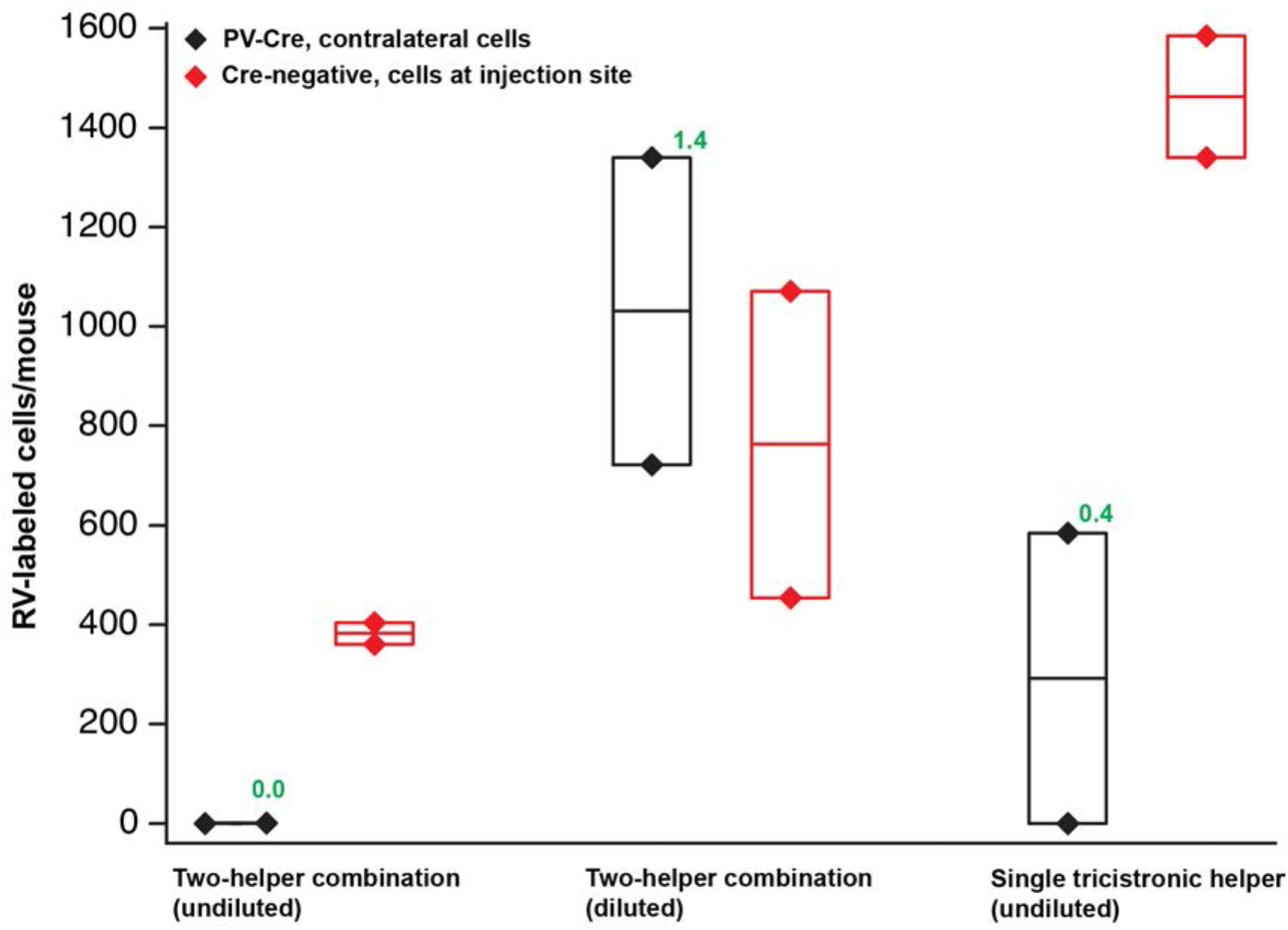
Quantification of results in PV-Cre and Cre-negative mice: pilot study with helper viruses either undiluted or diluted to titers used in previous work. Black depicts numbers of neurons labeled by RV in contralateral cortex (in every other 50 μm section) in PV-Cre mice; red depicts numbers of RV-labeled neurons in the vicinity of the injection site, i.e. in ipsilateral cortex (in every other 50 μm section) in Cre-negative mice. for all conditions. Diamonds represent cell counts from individual mice; the middle lines in the boxes represent the average count for each condition. Numbers in green represent the ratio of contralateral neurons in Cre+ mice to ipsilateral neurons in Cre-mice. “Diluted” here means diluted to the titers of other batches used previously in our laboratory; see main text for details. Excessive concentrations of helper viruses gave very poor results. Source numbers are provided in table S1.

We therefore embarked on a systematic set of experiments testing a range of dilutions for each helper virus, in order to find a set of dilutions for both the two-helper combination and the single helper that would result in efficient transsynaptic label in Cre mice but low background label in Cre-negative mice. For these experiments, the rabies virus used was RVΔG-4mCherry(EnvA)[16], expressing mCherry[34] rather than FLPo, to correspond most closely with the kind of experimental design used by typical users (the use of the FLP/FRT system in the pilot experiments described above was because those were originally intended to be controls for a different project). The results of these experiments are quantified in Figure 5, with source numbers given in table S2.

**Figure 5:**
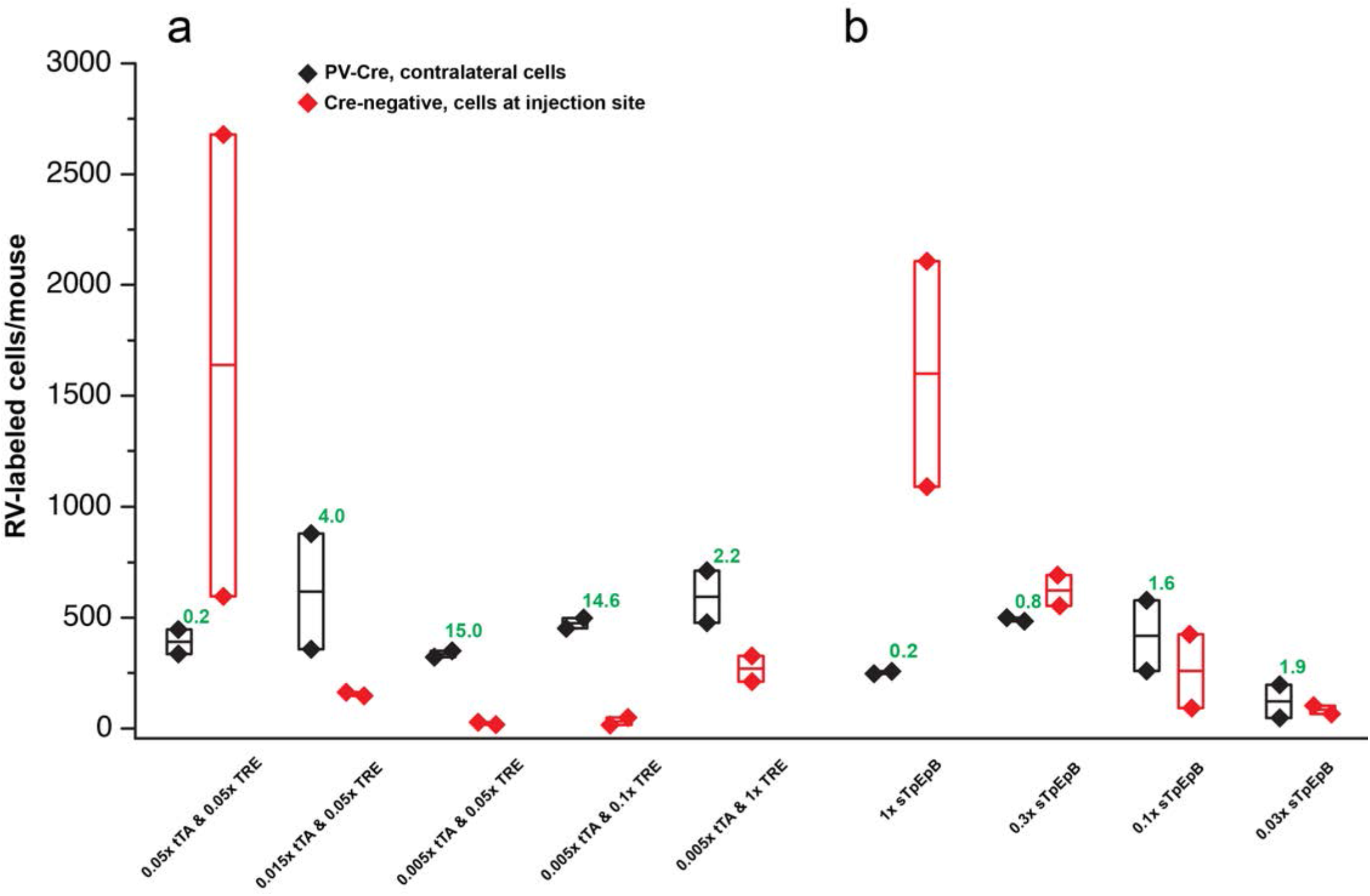
Quantification of results in PV-Cre and Cre-negative mice: systematic dilution series. a) Results of varying the concentrations of the two helper viruses in the tTA-TRE combination system. The highest ratio of contralateral neurons in Cre+ mice to ipsilateral neurons in Cre-mice was obtained with a 1:200 dilution of AAV1-syn-FLEX-splitTVA-EGFP-tTA and a 1:20 dilution of AAV1-TREtight-mTagBFP2-B19G (“0.005x tTA & 0.05x TRE” in the figure). b) Results of varying the concentration of the single helper virus AAV1-syn-FLEX-splitTVA-EGFP-B19G. Higher dilutions (out to 1:33.3) give higher ratios of contralateral neurons in Cre+ mice to ipsilateral neurons in Cre-mice, but results with the single-helper approach were nowhere near as good as with the two-helper combination. n=2 for all conditions. Diamonds represent cell counts from individual mice; the middle lines in the boxes represent the average count for each condition. Source numbers are provided in table S2.

For the two-helper combination (Figure 5, panel a), we were able to find dilutions that resulted in good transsynaptic label in Cre mice with little label at the injection site of Cre-negative mice. We began with 1:20 dilutions of both AAV1-syn-FLEX-splitTVA-EGFP-tTA and AAV1-TREtight-mTagBFP2-B19G, approximating (with simplification) the 1:17.96 & 1:19.98 dilutions used for used for Figures 2-4. Holding the concentration of the TRE AAV constant, we compared dilutions of the FLEX AAV of 1:20, 1:66.67, and 1:200 (labeled in panel a as “0.05x”, “0.015x”, and “0.005x”, respectively). Of these, we found that the most extreme dilution tested, 1:200, worked best, with the numbers of labeled cells in contralateral cortex in Cre mice almost as high as for the 1:20 dilution but with the numbers of labeled cells at the injection site in Cre-negative mice drastically reduced. Holding the concentration of the FLEX AAV constant at 1:200, we then tried increasing the concentration of the TRE AAV, comparing the 1:20 dilution to 1:10 and to undiluted stock. Interestingly, while these two additional conditions resulted in somewhat higher numbers of labeled contralateral cells in PV-Cre mice, they also greatly increased the numbers of cells labeled at the injection site in Cre-negative mice (compare “0.005x tTA & 0.05x TRE” to “0.005x tTA & 1x TRE” in Figure 5, panel a). Because the amount of TVA was not changed across these latter conditions, we assume that the additional labeled cells in Cre-negative mice were due to leaky G expression being sufficient to allow limited transsynaptic spread of the rabies virus from the initially-infected cells.

The best condition tested was therefore AAV1-syn-FLEX-splitTVA-EGFP-tTA 1:200 (for a final titer of 8.5 × 10^10 gc/ml) and AAV1-TREtight-mTagBFP2-B19G 1:20 (for a final titer of 1.6 × 10^12 GC/ml). Example images of results using this combination are shown in Figure 6, panels a-h for Cre mice and Figure 6, panels i-l for Cre-negative mice.

**Figure 6:**
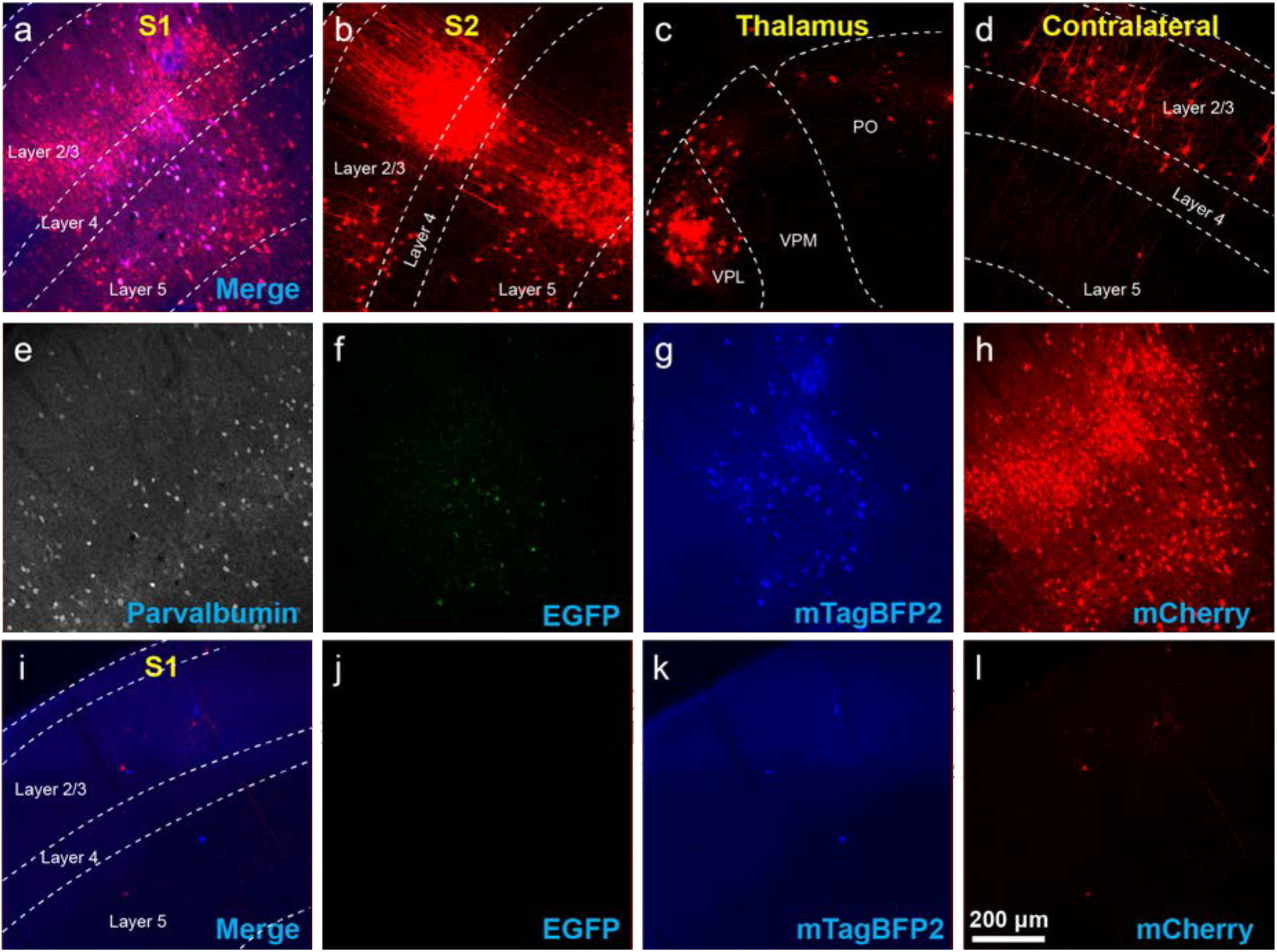
Example results using recommended dilutions of two-helper combination. a-h) Use of the two-AAV combination at 1:200 and 1:20 dilutions (see main text) labeled many presynaptic neurons. a) Injection site in S1. Green = anti-EGFP staining, blue = mTagBFP2, red = mCherry. Individual channels from this field are shown in panels e-h. b) Many labeled neurons were found in ipsilateral S2. c) Many labeled neurons were found in ipsilateral thalamus (VPL, VPM and Po). d) Many labeled neurons were found in contralateral S1. e-h) Individual channels from the field shown in panel a. e) anti-parvalbumin staining (not shown in panel a). f) anti-EGFP staining, indicating expression from the first, Cre-dependent AAV. Note that, at this dilution, EGFP signal is quite dim even with immunoamplification. g) mTagBFP2, indicating expression from the second, tTA-dependent AAV. h) mCherry, indicating presence of the ΔG rabies virus. i-l) Injection site after using two-helper combination at 1:200 & 1:20 (see main text): few mCherry-labeled cells are seen. i) Overlay of j-l. j) anti-EGFP staining: no signal is visible, even with amplification. k) mTagBFP2 signal. A few blue cells are seen even at these dilutions. l) mCherry expressed by RV. Scale bar in a: 200 μm, applies to all panels.

For the single tricistronic helper AAV1-syn-FLEX-splitTVA-EGFP-B19G, we compared undiluted (“1x” in Figure 5 panel b) to 1:3.33, 1:10, and 1:33.3 dilutions (“0.3x”, “0.1x”, and “0.03x” in the figure). While all the diluted versions improved matters over the undiluted version, none of the dilutions gave particularly good results, with even the highest dilution still resulting in much higher numbers of labeled cells at the injection site in Cre-negative animals than were found with the optimized two-helper combination, but with far fewer transsynaptically labeled cells in Cre mice. Example images of results using the 1:10 dilution are shown in Figure 7, panels a-g for Cre mice and Figure 7, panels h-j for Cre-negative mice.

**Figure 7:**
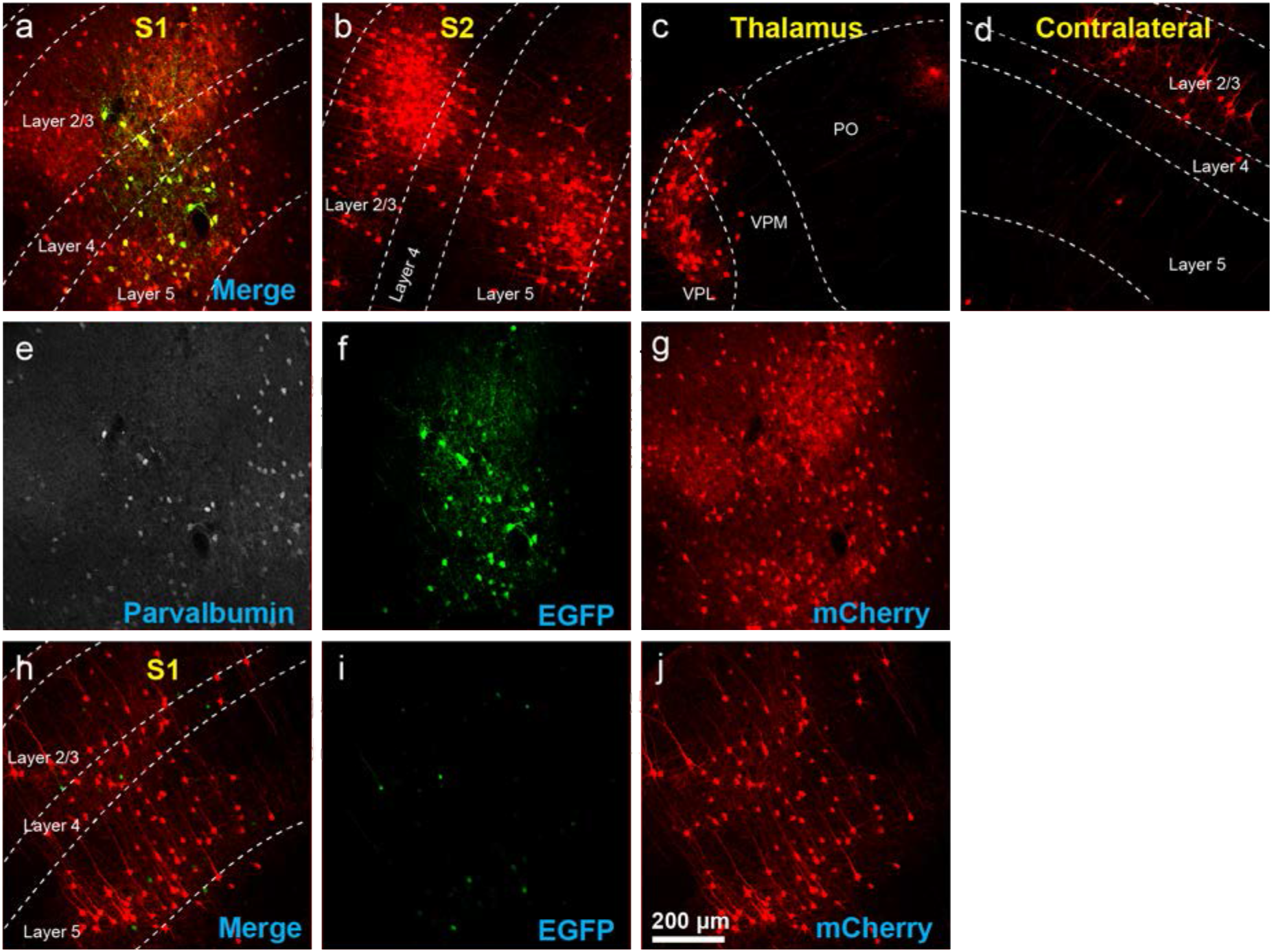
Results using recommended dilutions of the single tricistronic helper. a-g) Use of the single tricistronic helper AAV at 1:10 dilution (see main text) also labeled many presynaptic neurons (but see Figure 7). a) Injection site in S1; individual channels from this field are shown in panels e-g. b) Labeled neurons in ipsilateral S2. c) Labeled neurons in ipsilateral thalamus (VPL, VPM and Po). d) Labeled neurons in contralateral S1. e) anti-parvalbumin staining (not shown in panel a). f) anti-EGFP staining, indicating expression from the first, Cre-dependent AAV. g) mCherry, indicating presence of the ΔG rabies virus. h-j) Injection site after using single tricistronic helper: many mCherry-labeled neurons are present. h) Overlay of i-j. i) anti-EGFP staining: significant signal is seen even in these Cre-negative mice. j) mCherry expressed by RV. Scale bar in a: 200 μm, applies to all panels.

Finally, to determine whether the dilutions for the two-virus combination that worked best in PV-Cre mice also worked well in another Cre line, we performed a similar experiment in DAT-Cre mice[37], with 1:200 AAV1-syn-FLEX-splitTVA-EGFP-tTA and 1:20 AAV1-TREtight-mTagBFP2-B19G followed by RVΔG-4FLPo(EnvA) a week later. As shown in Figure 8 (panels a-h for Cre mice and panels i-l for Cre-negative mice), there were abundant labeled cells in striatum and cortex, suggesting that these helper virus dilutions may work well with other starting cell populations.

**Figure 8:**
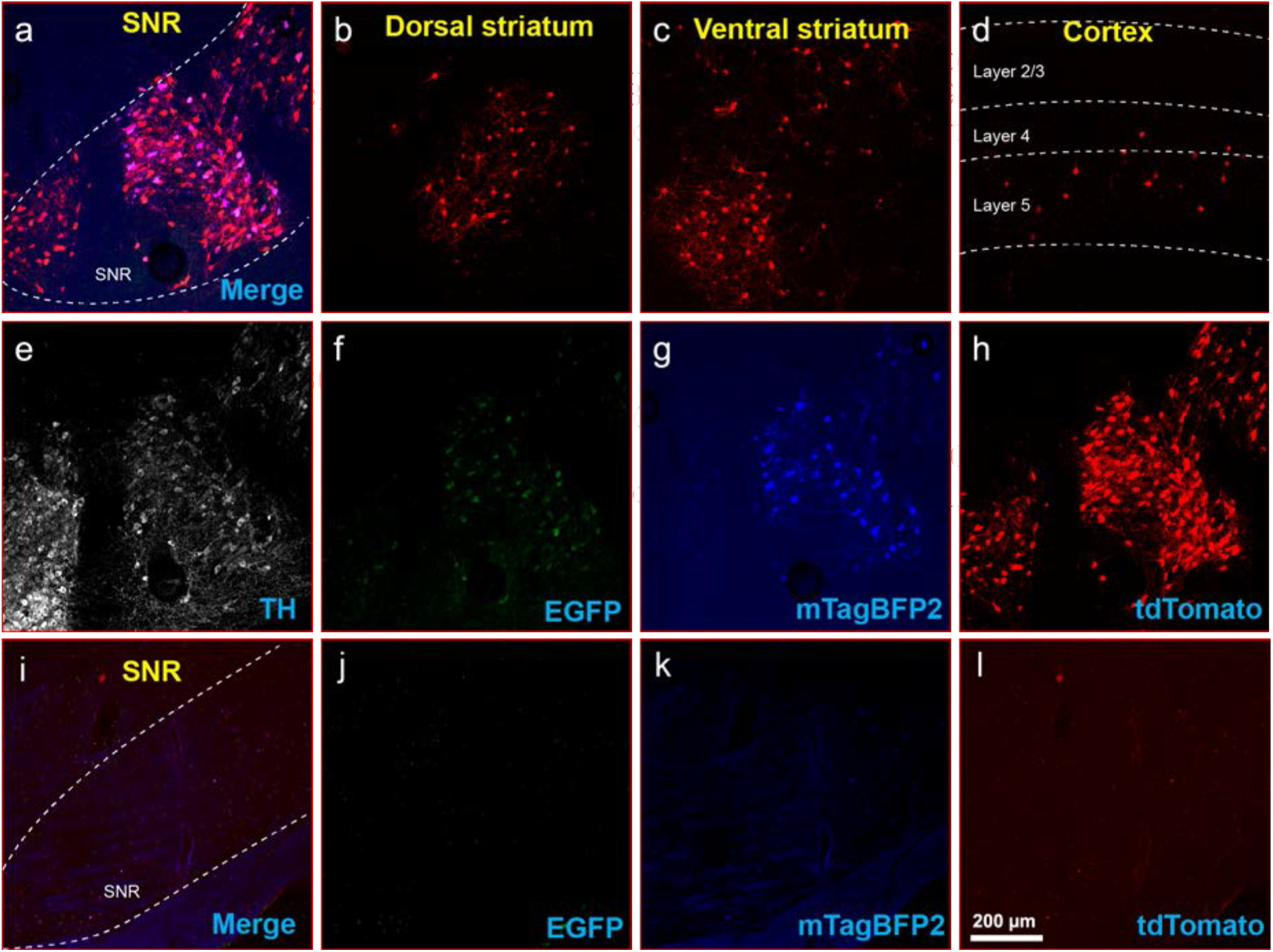
Inputs to midbrain dopaminergic cells using recommended dilutions of two-helper combination. a-h) Results in DAT-Cre mice. a) Injection site in substantial nigra reticulata (SNR): overlay of panels f-h. b) RV-labeled cells in dorsal striatum. c) RV-labeled cells in ventral striatum. d) RV-labeled cells in cortex. e) anti-tyrosine hydroxylase (TH) staining (not shown in panel a). f) anti-EGFP staining, indicating expression from the first, Cre-dependent AAV. g) mTagBFP2, indicating expression from the second, tTA-dependent AAV. h) tdTomato, reporting activity of the FLPo-encoding rabies virus. i-l) Results in Cre-negative mice (injection site shown). i) Overlay of j-l. j) anti-EGFP staining: no signal is visible, even with immunostaining. k) mTagBFP2 signal. no signal is visible, even with amplification. l) tdTomato, reporting activity of the FLPo-encoding rabies virus. Only one labeled cell is visible. Scale bar in a: 200 μm, applies to all panels.

## DISCUSSION

Our results demonstrate that the success of monosynaptic tracing depends strongly on the complementation strategy (using the two tTA-TRE coupled helper viruses worked much better than the single helper virus expressing TVA, EGFP, and G) and on the concentrations of the helper viruses.

More generally, our findings underscore that monosynaptic tracing results should not be taken as a complete delineation of the set of cells presynaptic to a targeted starting cell group, given that the number of false negatives (unlabeled cells that are actually presynaptic to the starting cells) clearly depends on the experimental parameters.

We have done these titration experiments with cortical injections in PV-Cre or Cre-negative mice. Results with other injection sites and Cre lines will presumably vary depending on the tropism of AAV1 for the targeted cell type and the other cells in the vicinity of the injection site. However, the fact that the parameters that we found to work best in PV-Cre, namely the two-helper combination with dilutions of 1:200 and 1:20, respectively, also gave good results in DAT-Cre mice (Figure 8) may indicate that these could be good general-purpose parameters for most Cre lines; at the very least, they should serve as a good starting point for a much more limited set of titration experiments than we have undertaken here.

## METHODS

All experiments involving animals were conducted according to NIH guidelines and approved by the MIT Committee for Animal Care. Mice were housed 1-5per cage under a normal light/dark cycle for all experiments.

### Viruses

Cloning of AAV genome plasmids pAAV-synP-FLEX-splitTVA-EGFP-B19G (Addgene 52473), pAAV-syn-FLEX-splitTVA-EGFP-tTA (Addgene 100798), and pAAV-TREtight-mTagBFP2-B19G (Addgene 100799) has been described[16; 18]. These genomes were packaged in serotype 1 AAV capsids by, and are available for purchase from, Addgene (catalog numbers 52473-AAV1, 100798-AAV1, and 100799-AAV1). The titers of the AAVs, as determined by Addgene by qPCR, were as follows:

AAV1-synP-FLEX-splitTVA-EGFP-B19G (lot # v14715): 2.4 × 10^13 gc⁄ml
AAV1-syn-FLEX-splitTVA-EGFP-tTA (lot # v15287): 1.7 × 10^13 gc⁄ml
AAV1-TREtight-mTagBFP2-B19G (lot # v14716): 3.2 × 10^13 gc⁄ml.

Cloning of pRVΔG-4FLPo[38] (Addgene 122050) and pRVΔG-4mCherry[39] (Addgene 52488) have been described. Production of EnvA-enveloped rabies virus RVΔG-4mCherry(EnvA)[16] was done as described previously[40; 41; 42] but using helper plasmids pCAG-B19N (Addgene #59924), pCAG-B19P (Addgene #59925), pCAG-B19G (Addgene #59921), pCAG-B19L (Addgene #59922), and pCAG-T7pol (Addgene #59926) for the rescue step[42]. The final titers were 5.82 × 10^9 infectious units/ml for RVΔG-4FLPo(EnvA) and 1.70 × 10^10 infectious units/ml for RVΔG-4mCherry(EnvA), as determined by infection of TVA-expressing cells as described previously[38; 40].

Helper AAVs were diluted in Dulbecco’s phosphate-buffered saline (DPBS) (Fisher, 14-190-250) by the desired dilution factors (see main text). In the case of AAV-syn-FLEX-splitTVA-EGFP-tTA and AAV-TREtight-mTagBFP2-B19G, the two viruses were combined (after dilution, when applicable) in a 50/50 ratio by volume before injection.

### Mouse strains

Adult mice of both sexes were used. For compatibility with other projects in the laboratory, the PV-Cre (Jackson 017320) and DAT-Cre (Jackson 006660) used were also heterozygous for the FLP-dependent tdTomato reporter line Ai65F[35](obtained in this case by crossing the Cre- and FLP-dependent tdTomato double-reporter line Ai65D[43] (Jackson Laboratory 021875) to the Cre deleter line Meox2-Cre[44] (Jackson Laboratory 003755), then breeding out the Meox2-Cre allele, resulting in a reporter line for which only FLP is required for expression of tdTomato). For those mice in which RVΔG-4FLPo was used, the reporter allele was necessary for reporting RV activity; for those in which RVΔG-4mCherry was used, the presence of this reporter allele was irrelevant. For Cre-negative control injections using RVΔG-4FLPo, the Ai65F line was used. For Cre-negative control injections using RVΔG-4mCherry, either Ai65F or the Cre-dependent reporter line Ai14[45] was used; in these cases, the presence of the reporter alleles was again irrelevant.

### Stereotaxic injections

For pilot studies (Figures 2-4), we injected 300 nl of helper AAV solution into primary somatosensory cortex (coordinates with respect to bregma: APX= −0.58 mm, LM = 3.00 mm, DV = −1.00 mm) of anesthetized adult mice as described[32], using a stereotaxic instrument (Stoelting Co., 51925) and custom injection apparatus consisting of a hydraulic manipulator (Narishige, MO-10) with headstage coupled via custom adaptors to a wire plunger advanced through pulled glass capillaries (Drummond, Wiretrol II) back-filled with mineral oil and front-filled with viral vector solution. 7 days after AAV injection, 100 nl of RVΔG-FLPo(EnvA) was injected in the same site.

For subsequent experiments in PV-Cre (Figures 5-7), 200 nl of helper AAV solution was injected, followed by 100 nl of RVΔG-4mCherry(EnvA) 7 days later. For DAT-Cre mice (Figure 8), 200 nl of helper AAV solution was injected (AP= −3.00 mm, LM = 1.50 mm, DV = −4.20 mm), followed by 500 nl of RVΔG-FLPo(EnvA) 7 days later. Two mice were used for each condition (n=2).

### Perfusions and histology

7 days after injection of rabies virus, mice were transcardially perfused with 4% paraformaldehyde in phosphate-buffered saline. Brains were postfixed in 4% paraformaldehyde overnight on a shaker at 4°C and cut into 50 μm coronal sections on a vibrating microtome (Leica, VT-1000S), with sections collected into six tubes (containing cryoprotectant solution as described[32]) each, so that each tube contained a series of every sixth section through the sectioned region of the brain. For confocal imaging, sections were immunostained as described previously [45] with the following antibodies (as applicable) at the following respective dilutions: chicken anti-GFP (Aves Labs GFP-1020) 1:1000, guinea pig anti-parvalbumin (Synaptic Systems 195004) 1:1000, sheep anti-tyrosine hydroxylase (Millipore AB1542)) 1:1000, with secondary antibodies donkey anti-chicken Alexa Fluor 488 (Jackson Immuno 703-545-155) 1:200, donkey anti-guinea pig, AlexaFluor 647 conjugated (Jackson Immuno 706-605-148) 1:200, and donkey anti-sheep, AlexaFluor 647 conjugated (Jackson Immuno 713-605-147)) 1:200. Sections were mounted with Prolong Diamond Antifade mounting medium (Thermo Fisher P36970) and imaged on a confocal microscope (Zeiss, LSM 710). For counts, every other series (i.e., either series 1, 3, and 5 or series 2, 4, and 6) of each brain was mounted, so that 50% of the sections from each brain were mounted and examined.

### Counts

Neurons labeled with either tdTomato or mCherry in contralateral cortex in PV-Cre mice (crossed with Ai65F reporter mice; see Mouse Strains) and at the injection site in Cre-negative (Ai65F reporter) mice were counted manually by examining every other 50 μm section (i.e., 50% of the sections) on an epifluorescence microscope (Zeiss, Imager.Z2). Counts of cells in the contralateral cortex were restricted to those in the sectioned anterior-posterior region common to all sectioned brains. This encompassed the sections between 1.2 mm and −3.0 mm relative to bregma[46].

## ACKNOWLEDGEMENTS

We thank Kimberly Ritola of Janelia Research Campus for helpful discussions regarding the important issue of leaky FLEX viruses as well as Melina Fan and Karen Guerin of Addgene for helpful discussions and for producing the helper AAVs and providing them to our laboratory and to the worldwide neuroscience community. Research reported in this publication was supported by BRAIN Initiative awards U01MH106018, U01MH114829, and U19MH114830 from the National Institute of Mental Health. This manuscript has been released as a preprint at bioRxiv[47].

---

**Figure S1:**
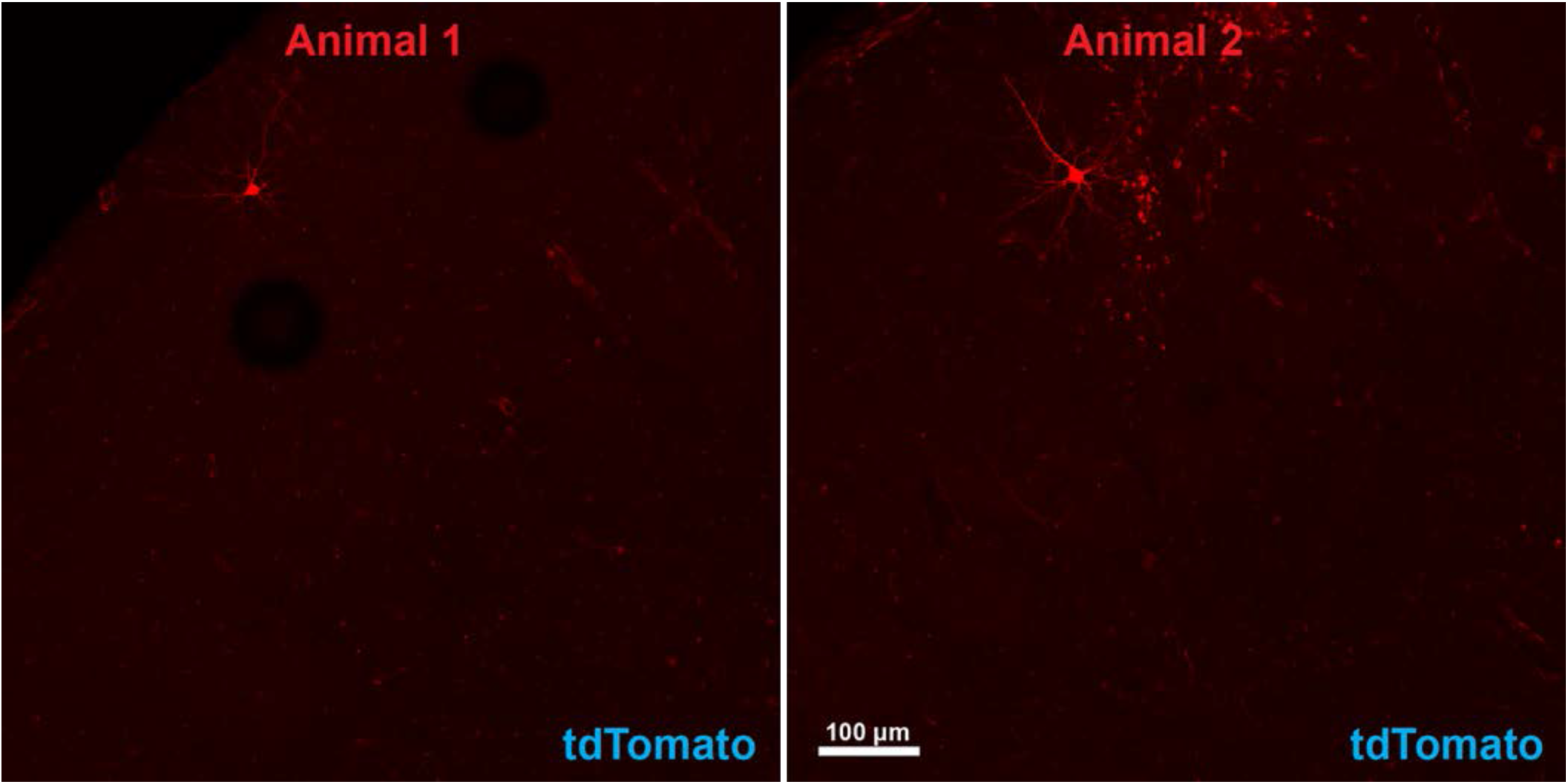
Results with rabies virus injection only without helper virus. S1 cortex of two animals were injected with 100 nl RVΔG-4FLPo(EnvA) virus only, without a previous AAV injection. Very few tdTomato-labelled cells were found. Scale bar in a: 100 μm, applies to all panels.

**Table S1 and S2:**
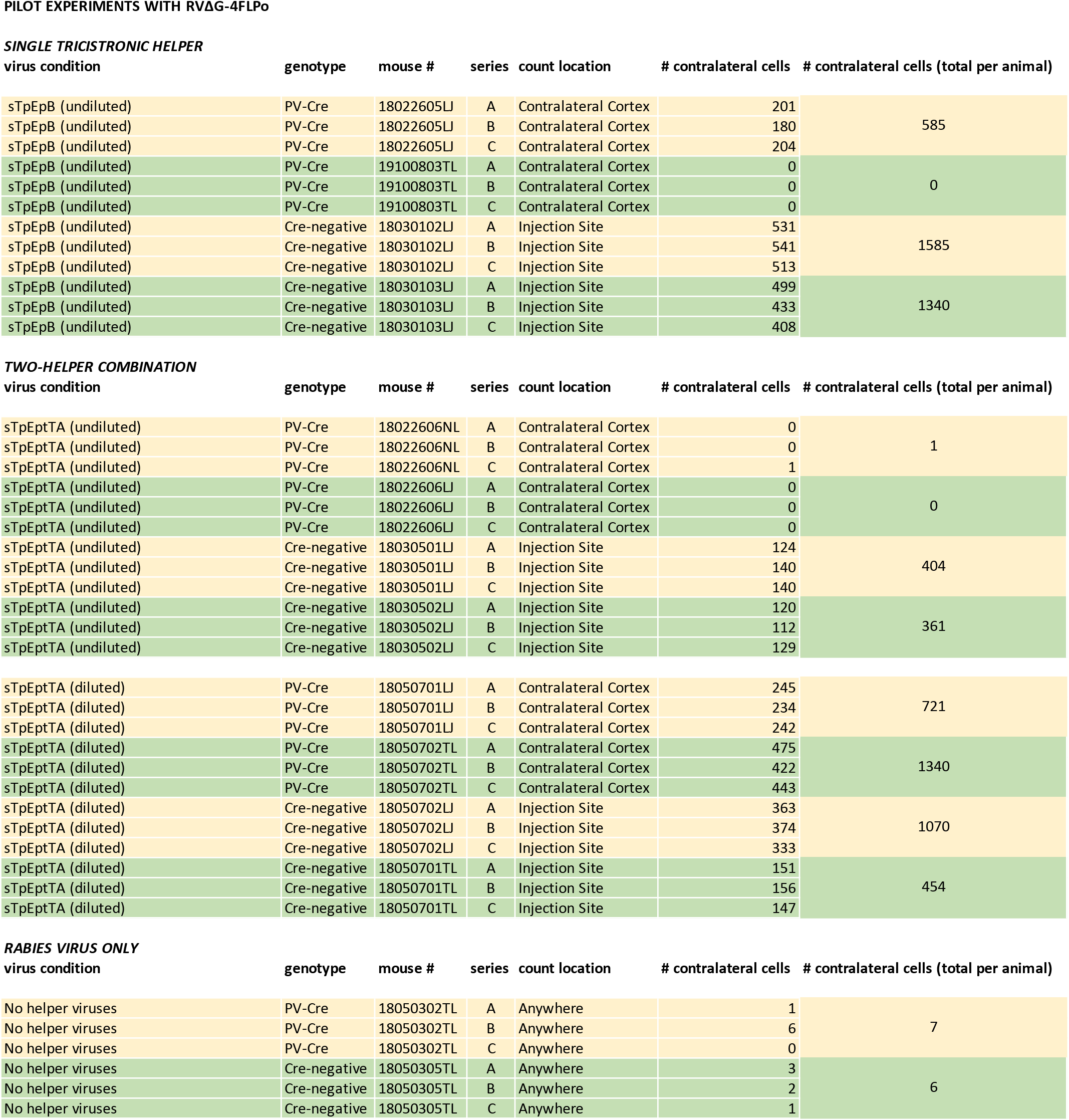
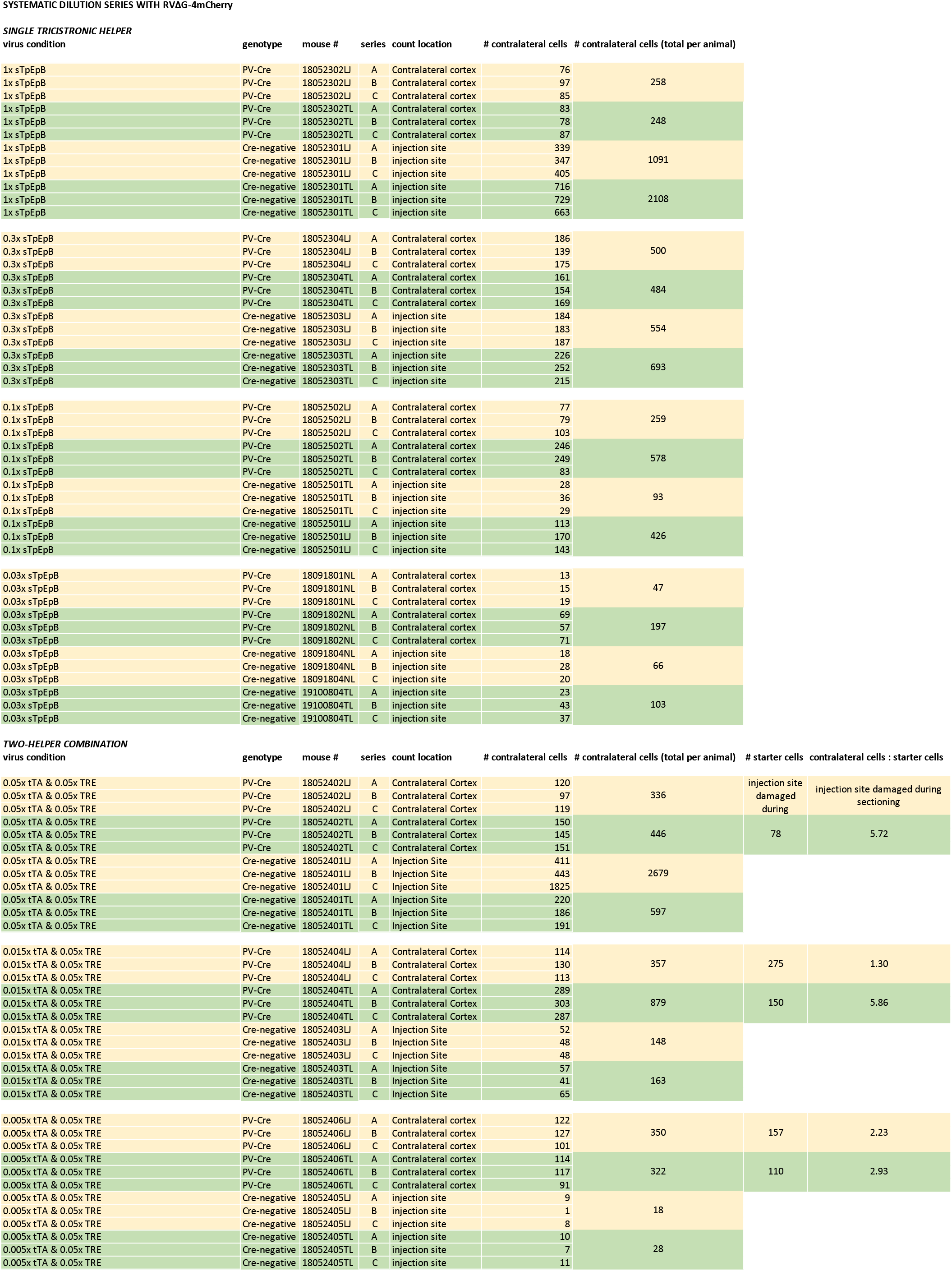
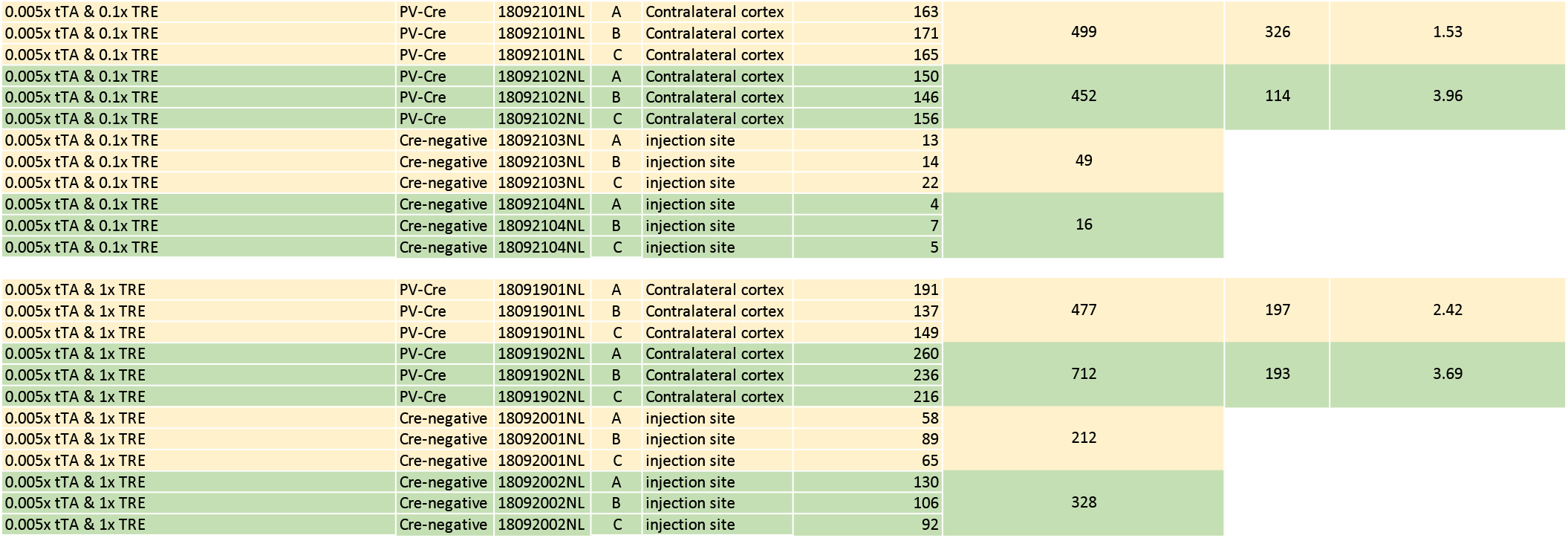
Counts of labeled cells in PV-Cre and Cre-negative mice. Numbers of labeled neurons in contralateral cortex of PV-Cre mice and at the injection site of Cre-negative mice for the various helper virus conditions. Each number in the “# cells” column indicates the total number of labeled cells found in the examined region (either the vicinity of the injection site or the contralateral cortex) across all 50 μm sections in that series of every sixth section (see methods). The total number of labeled neurons counted for a given mouse was the sum of the total labeled neurons in each of the three examined series for that mouse (i.e., the total found in every other section). The means of the total numbers of labeled neurons and individual count for each condition are graphed in Figures 4 (Table S1) and 5 (Table S2).

